# *Withania somnifera* root extract reduces susceptibility of the model worm *Caenorhabditis elegans* to different types of stressors

**DOI:** 10.64898/2025.12.18.695325

**Authors:** Nidhi Thakkar, Dilip Mehta, Vijay Kothari

## Abstract

**Background:** *Withania somnifera*, commonly known as *Ashwagandha*, is one of the most widely prescribed plants in the Indian system of traditional medicine. Its roots are believed to impart comprehensive health benefits supporting lifespan and healthspan through its anti-stress, immunomodulatory, and neuroprotective activity. Scientific validation of its claimed biological effects is warranted.

**Methods:** The nematode worm *Caenorhabditis elegans* was challenged with heat or certain neurotoxic/mitotoxic chemical stressors, and worm activity (healthspan) was compared between *W. somnifera* root extract (WSRE)-exposed and control worm populations, through an automated worm tracker.

**Results:** Previous exposure of worms to WSRE conferred prophylactic benefit on them in face of subsequent challenge with rotenone, MnCl_2_, or levamisole. Worms pre-exposed to either of these toxins were able to recover faster in presence of WSRE. This extract was also able to reduce the susceptibility of wild type worms to heat-induced paralysis, and that of transgenic worms to beta-amyloid mediated paralysis.

**Conclusions:** Adaptogenic potential of *W. somnifera* roots was demonstrated in worm model. WSRE can be said to possess a wide anti-stress spectrum as it protected the worms against different stressors inducing toxicity through different mechanisms including interference with acetylcholine and dopamine neurons and function. Our results support the candidature of *W. somnifera* roots as a potent nutraceutical.

## 1. Introduction

Stress is an integral part of life, and to combat various stress types, life forms have evolved different types of stress responses. While certain stressors have an immediate short-term effect, other may have chronic effect influencing quality of life over longer period. While different stressors have different modes of action, certain effects are common in the target biological system, e.g. build-up of reactive oxygen species, mitochondrial dysfunction, etc. Latter is commonly associated with neurodegenerative diseases (Johri and Beal, 2012) characterized by the progressive loss of neuronal structure and function. Neurodegenerative diseases pose a significant challenge to public health worldwide, as they directly are associated with deteriorated quality of life, particularly in aged people. Since humans come in contact of a diverse variety of stressors of physicochemical or biological origin, finding some broad-spectrum therapeutic formulation, which can protect against multiple stressors can be an attractive strategy. Developing therapeutic strategies to mitigate the health-deteriorating effects of different stress types (e.g. toxic chemicals, heat, etc.) remains a critical area of research.

Many plants in traditional medicine have been mentioned to possess stress-alleviating property. One such plant with strong standing in the Indian system of Medicine (*Ayurved*) is *Withania somnifera* (L.) Dunal, commonly known as *Ashwagandha*. In ancient medical literature, this plant is recognized as a ‘Rasayana’, which means an adaptogen that can help the body cope with stress, anxiety and fatigue. Modern literature has also indicated *W. somnifera* roots to possess immunomodulatory, neuroprotective, and anti-depressant activities (Vaidya et al., 2024). It is considered as a potent nutraceutical (Saha et al., 2024). Though *W. somnifera* is a relatively well-researched plant, addition of more evidence regarding its safety and beneficial effects to the existing body of literature can help build higher public confidence in it. At times, either due to adulteration or due to deviation from the traditionally prescribed method of use of a particular plant, regulatory agencies and public may grow suspicious even of the plants with a long history of safe and efficacious usage. Such situations can best be handled through rigorous, transparent research, and informed regulations to address safety concerns while preserving the credibility and global acceptance of widely used *ayurvedic* herbs like *W. somnifera*. Mainstreaming the use of any herbal preparation for therapeutic or nutraceutical applications relies on demonstration of claimed biological activities in them through robust scientific assays (Verpoorte, 2017).

Earlier we had reported beneficial effect of *W. somnifera* root extract on lifespan, healthspan, and fertility of the model worm *Caenorhabditis elegans* (Thakkar et al., 2025). Now we report its ability to protect this model worm upon being challenged with heat or toxic chemicals. This nematode worm in recent years has proved to be a useful model for stress biology research owing to its conserved stress response pathways, and its ability to adapt to different stressors, through epigenetic changes that offer protection against age-associated decline (Deji-Oloruntoba et al., 2025). Confidence of the research community in relevance of this worm model for neurotoxicity testing is also increasing (Sammi et al., 2022).

## 2. Methods

### 2.1. Plant extract

The hydroalcoholic extract (LongeFera™; Batch no. PHPL/RD/114/EXT-094-01) of *W. somnifera* root was supplied by Phytoveda Pvt. Ltd., Mumbai. Background details on collection and authentication of the plant material, extract preparation, etc. are available in our previous study (Thakkar et al., 2025) on the same extract. This extract meets the relevant quality control requirements of the United States Pharmacopeia (USP). Total withanolides content in this extract was determined to be 2.69 ± 0.02%. Chromatographic fingerprint of this extract, along with the concentration of major phytocompounds like withanosides and withanolides can be viewed as Figure S1 at: https://journals.aboutscience.eu/index.php/dti/article/view/3368/3962.

For bioassay purpose, four gram of the extract powder was suspended in 10 mL of sterile distilled water, and kept for shaking at ambient temperature for 30 min. The insoluble fraction from the aqueous suspension was removed by centrifugation (7500 g; 25°C; 10 min). The soluble fraction (supernatant) was passed through a 0.45 µm syringe filter (Axiva), and the filtrate was stored in a sterile glass vial (15 mL; Borosil) under refrigeration. Solubility of the extract in water was calculated to be 69.77%.

### 2.2. Test organism

Except for the thermorecovery assay, all assays were performed with the wild type N2 Bristol strain of *C. elegans* procured from the Caenorhabditis Genetics Center (Minneapolis, USA). Strain CL4176, used for the thermorecovery assay, was received as a gift from the Xavier’s Research Foundation, Ahmedabad. This strain produces beta amyloid peptide (https://cgc.umn.edu/strain/CL4176) when incubation temperature is upshifted from 15°C to 35°C, and is relevant to research on diseases involving neurodegeneration (Xu et al., 2025).

Lyophilized *E. coli* OP50 (Biovirid, Netherlands) was used as food for *C. elegans*, while maintaining the worm on NGM agar plates (Nematode Growing Medium). Worm synchronization was done as described in literature (Corsi et al., 2015). Prior to the *in vivo* assays, worms were kept without food for two days to make them gnotobiotic.

### 2.3. Chemical stressors

Chemical stressor used in this study included rotenone (Merck; R8875), levamisole (Merck; T1512-2G), nonanol (Merck; 157471000), ivermectin (Merck; I8898), benzimidazole (HiMedia; GRM1105-25G), aluminum chloride (027073), manganese chloride (HiMedia; RM 685-500G), paraquat (Merck; 1910-42-5), and hydrogen paroxide (Merck; 1.93408.0521). H_2_O_2_ or parquet were dissolved in water whereas rotenone, levamisole, ivermectin or benzimidazole were dissolved in DMSO (Merck). Nonanol was added directly into the assay wells containing worms in M9 buffer. Since AlCl_3_ and MnCl_2_ were getting precipitated in M9 buffer, for assays involving them, worms were kept in distilled water. Additional details on all these stressors are provided in Table-1.

**Table 1.** List of chemical stressors used in this study.

| Sr. No. | Name of Chemical | Type of toxin | Effect on the worm | Additional Remarks | Reference |
| --- | --- | --- | --- | --- | --- |
| 1 | Rotenone | Mitotoxic and neurotoxic | Inhibits mitochondrial complex I, causes ATP depletion, and dopaminergic neuron degeneration | Widely used to study the effects of mitochondrial dysfunction on dopaminergic neurons in the context of Parkinson's disease. | Gonzalez-Hunt et al., 2021 |
| 2 | Paraquat | Mitotoxic | Induces mitochondrial dysfunction, and dopaminergic neurodegeneration | Exposure to paraquat in humans is associated with a higher risk of developing neurological diseases such as Parkinson's. | Bora et al., 2021 |
| 3 | Aluminum chloride (AlCl<sub>3</sub>) | Neurotoxic | Impairs neurotransmitter synthesis, induces apoptotic neuronal death, and disrupts central nervous system homeostasis | Al exposure in humans is suggested to be linked with dementia, osteomalacia and microcytic anaemia. | Page et al., 2012 |
| 4 | Manganese Chloride (MnCl<sub>2</sub>) | Neurotoxic and mitotoxic | Triggers dopaminergic neuron degeneration, and mitochondrial dysfunction | Accumulation of Mn in the brain results in Manganism that presents with Parkinson's disease (PD)-like symptoms. | Moline, 2023; Chen et al., 2015 |
| 5 | Ivermectin | Anthelmintic | Affects ligand-gated chloride channels | Useful for nematode motility studies | Hernando and Bouzat, 2014; Patil et al., 2022 |
| 6 | Benzimidazole | Anthelmintic | Disrupts microtubules in neurons, disrupts locomotion and reproduction; detrimental effect on oocytes; Disrupts processes requiring integral microtubules | Used as positive control in anti-nematode assays | Holden-Dye and Walker, 2014 |
| 7 | Nonanol | Neurotoxin | Triggers aversive olfactory behaviour | Used in assays for indirect assessment of dopamine levels | Sammi et al., 2022 |
| 8 | Levamisole | Neurotoxic | Targets nicotinic acetylcholine receptors, causing neuromuscular paralysis | Used to treat roundworm infections in humans and animals | Martin et al., 2012 |
| 9 | Hydrogen peroxide | Oxidant | Generates oxidative stress | Widely used to induce oxidative damage in cells or animal models | Fu et al., 2015 |

### 2.4. Toxicity assays

#### 2.4.1. Prophylactic assay

To assess the anti-stress or toxicity-alleviating potential of the *W. somnifera* root extract, worms pre-incubated with the test extract (600 ppm) were subsequently challenged with various stressors (Table-1), and their survival was compared with that of control population (challenged with stressors without any extract pre-exposure). While worm survival, morphology, and paralysis (if) induced by stress were observed microscopically, overall activity at the population level was quantified through an automated worm tracker (WMicrotracker ARENA; Phylumtech, Argentina). Worm tracker used for capturing worm activity was set at 23°C, and acquisition lapse time was kept 15 min.

Different types of stressors (e.g. mitotoxins, neurotoxins, oxidizing agents, heat, etc.), at sub-lethal doses were used to induce stress in this worm. Appropriate dose(s) for each of the stressor was determined beforehand by challenging the worms with the stressor over a broad concentration range (Figure S1). We deliberately chose to use the extract-pre-treated worms for these assays in order to avoid simultaneous presence of toxin and the test extract in the assay well. This guards against complications in result interpretation owing to possible interaction of the extract with the stressor chemical. Concentration of the extract employed in this study was guided by our previous study (Thakkar et al., 2025), wherein the test extract was found to exert the optimum beneficial effect on worm healthspan and lifespan at 600 ppm.

Gnotobiotic worms were separated into two groups, control and experimental. The experimental worm population was incubated in 5 mL of M9 buffer supplemented with *W. somnifera* root extract (600 ppm final concentration) for 48 h at 22ºC, while the control worm group was kept in liquid media devoid of test formulation. The whole assay content was housed in conical flasks (25 mL capacity). Intermittent shaking was provided throughout the incubation for the sake of homogeneity, and to avoid settling of worms at the bottom. After incubation, content from each flask was transferred into sterile centrifuge tubes (15 mL) and rotated at 1000 rpm for 2 min. Resulting pellet was mixed with fresh sterile M9 buffer (5 mL), and again centrifuged. This washing step was done twice to ensure removal of any surface-attached residual extract from the experimental worm population. Worms obtained at the end of above processing were distributed in 24-well plates (HiMedia, Mumbai). Approximately, 100 worms (L3-L4) were added per well. Then, sub-lethal concentration of the stressor chemical was added, and the plates were incubated at 22ºC for five days. DMSO (0.5%v/v) was employed as a vehicle control, wherever applicable. Wells containing extract-pre-treated worms, not exposed to any toxin, were also included in the assays.

#### 2.4.2. Recovery assay

Once the toxins were identified, to which worm susceptibility was reduced following previous exposure to *W. somnifera* extract, with these toxins only, we conducted an additional assay, wherein toxin-pre-exposed worms were subsequently transferred into extract-supplemented media to check whether this extract can support better recovery of worms from the toxic shock. Gnotobiotic worms were separated into two groups, control and experimental. The experimental worm population was incubated in 5 mL of M9 buffer supplemented with any one toxin for 3-5 h at 22ºC, while the control worm group was kept in liquid media devoid of toxin. The whole assay content was housed in conical flasks (25 mL capacity). Intermittent shaking was provided throughout the incubation for the sake of homogeneity, and to avoid settling of worms at the bottom. After incubation, content from each flask was transferred into sterile centrifuge tubes (15 mL) and rotated at 1000 rpm for 2 min. Resulting pellet was mixed with fresh sterile M9 buffer (5 mL), and again centrifuged. This washing step was done twice to ensure removal of any surface-attached toxin from the experimental worm population. Worms obtained at the end of above processing were distributed in 24-well plates (HiMedia, Mumbai). Approximately, 100 worms (L3-L4) were added per well. Then, *W. somnifera* root extract (600 ppm final concentration) was added, and the plates were incubated at 22ºC for five days. Wells containing toxin-pre-treated worms, subsequently not exposed to *W. somnifera*, were also included in the assay.

Since a single positive control compound can not be recommended while working with different kinds of stressors, we employed multiple compounds as positive control, which have been reported in literature for their adaptogenic potential. They included cortisol (Yasuda et al., 2021) (2 ppm; Merck), dopamine (Chou et al., 2022) (5 mM; Samarth life sciences), ascorbic acid/vitamin C (Harrington and Harley, 1988) (250 µg/mL; Himedia), and caffeine (Dostal et al., 2010) (300 ppm; Merck).

### 2.5 Thermorecovery assay

For thermorecovery assay with wild type *C. elegans*, approximately 100 worms (L3-L4) were added per well in a 24-well plate. Each well contained 1 mL of M9 media with or without extract (600 ppm). These plates were kept first at 37°C for 3 h to give heat shock, read in the worm tracker, and then transferred into a 22°C incubator for five days. Qualitative microscopic observation and automated motility quantification was done on daily basis.

Thermorecovery assay was also done with the transgenic worm strain CL4176, which expresses human amyloid-beta protein in muscle cells, making it particularly sensitive to heat-induced paralysis leading to death. Ten gnotobiotic worms were placed on NGM agar supplemented with *W. somnifera* extract (600 ppm), in 35 mm glass dishes. Soon after placing the worms on agar, they were subjected to thermal stress by transferring the agar plates to an incubator set at 35°C for 4 h, followed by a recovery period at 22°C for 24 h. Control worms were treated in the same manner except that the agar used for them did not contain the plant extract. Worm survival was assessed microscopically by monitoring responses to mechanical stimuli, and the non-responsive worms were recorded as dead.

### 2.6 Metabolic activity assay

Alamar Blue^®^ assay was used to quantify the viability or metabolic activity of the worms. One hundred µL of Alamar Blue^®^ (Thermofisher) was added into each well containing approximately 100 worms in 900 µL of M9 media (with or without appropriate concentration of toxin or extract), making the total volume 1 mL, before incubation started. To quantify the amount of dye reduced, on last day of the experiment, content from wells was transferred into a separate plastic vial (1.5 mL), followed by centrifugation (13,600 g at 25°C) for 10 min. Then, the supernatant was read at 570 nm (Agilent Cary 60 UV-vis). Appropriate abiotic controls (containing the dye and other media components but no worms) were also included in the assay.

### 2.7 Statistics

All values reported (Mean ± SEM) are derived from three or more independent experiments, wherein each experiment contained three replicates (unless specified otherwise). Statistical significance was assessed using a t-test performed in Microsoft Excel^®^ (Version 2016), and data with p ≤ 0.05 were considered to be statistically significant.

## 3. Results and Discussion

Initially we challenged the extract-pre-exposed worms with nine different toxins (Table-1), and identified three (rotenone, MnCl_2_, and levamisole) of them to whom worm susceptibility was reduced owing to previous extract exposure. Another assay, wherein toxin-pre-exposed worms were subsequently incubated with the extract to investigate whether extract can support faster recovery from the toxic shock, was done only with those three toxins against which extract was found in first assay (prophylactic assay) to be effective. Results of both these assays with abovementioned three toxins are presented in following text, while results with those toxins against which extract did not offer any protection to the worms are presented in Figure S2.

### 3.1. *W. somnifera* reduces worm’s susceptibility to rotenone

Worms pre-fed with WSRE experienced a delayed death when challenged with rotenone (Figure 1A). While control worms experienced a 2.73-fold (p ≤ 0.001) reduction in their activity within first fifteen minutes of rotenone exposure, extract-fed worms exhibited activity, statistically at par, to the healthy worms (exposed neither to rotenone nor extract) i.e. 2.04-fold (p=0.001) higher than the rotenone-exposed worms. By the end of second day, when rotenone caused 89% (9.32-fold; p ≤ 0.001) reduction in worm activity in control population, extract-fed worms registered 4.77-fold (p=0.004) higher activity than rotenone-exposed worms. Though till last day of the experiment, extract-fed worms maintained higher activity, the absolute activity counts were quite low. On all days, dopamine-pre-fed worms showed activity higher than control worms in face of rotenone challenge.

**Figure 1.**
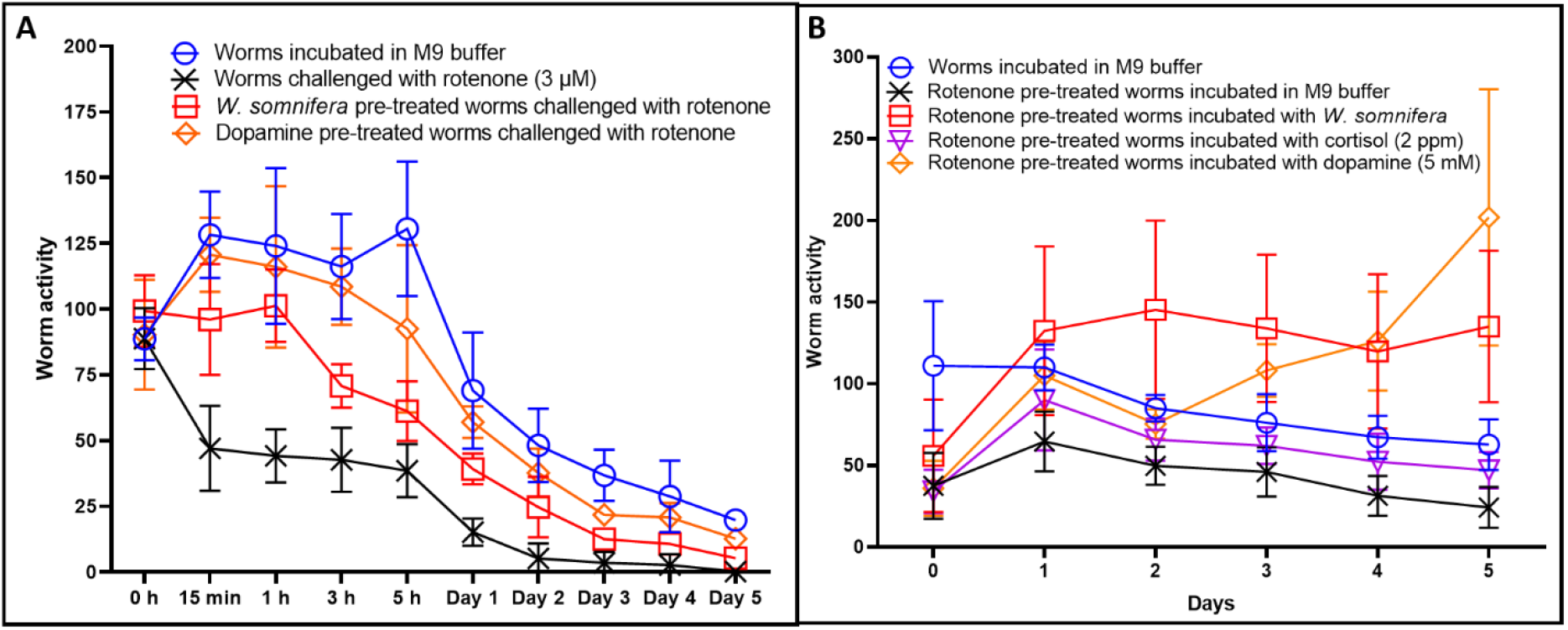
Protective effect of *Withania somnifera* against rotenone. **(A) Prophylactic effect**. Worms were pre-fed for 48 h with WSRE, ascorbic acid, cortisol, or dopamine, followed by continuous exposure to rotenone. Pre-feeding with either dopamine or WSRE noticeably reduced worm susceptibility to rotenone, supporting higher worm activity compared to the rotenone control. See Supplementary Videos A1-A2 and B1-B3. **(B) Recovery from toxic shock**. Worms were first exposed to rotenone for 5 h, and subsequently allowed to recover in presence of either the *W. somnifera* extract or positive control compounds. WSRE and dopamine, both supported worm recovery from toxic shock, and also supported progeny production on day four, whereas cortisol treatment resulted in a moderate increase in activity, lower than that observed with WSRE or dopamine. See Supplementary Videos A3-A4 and C1-C5. For clarity of presentation, lines pertaining to compounds (cortisol or ascorbic acid) showing no activity, which were running parallel to the rotenone control, are not shown in these graphs, so as to reduce visual crowding. WSRE: *Withania somnifera* root extract.

When toxin-pre-exposed worms were allowed to recover in absence or presence of WSRE, the worm population in presence of extract not only could recover faster from the toxic shock, but also was able to display fertility. It should be noted that presence of progenies does make a notable contribution to the worm activity count. By end of the fourth day, when toxin-pre-exposed worms showed almost half (53%) activity of that of control worms, those in wells pertaining to WSRE or dopamine displayed better activity (Figure 1B), bigger morphology as well as active reproduction. Cortisol was also able to support marginal recovery benefit to rotenone-pre-exposed worms. Till day four, extract’s effect was at par to that of dopamine, and particularly on day-2 WSRE supported worm activity 1.93-fold (p ≤ 0.001) higher than that supported by dopamine.

Since rotenone is known to be a mitotoxin, we hypothesized that the protective effect of WSRE against rotenone can emerge partly from its mitochondria-protective effect. As metabolic activity captured by quantifying reduction of certain dyes is a surrogate for mitochondrial activity (Springer et al., 1998), we quantified the metabolic activity/viability of worm population, incubated in presence or absence of WSRE followed by subsequent incubation in rotenone-supplemented media. Higher metabolic activity in WSRE-pre-fed worms facing rotenone challenge confirmed beneficial effect of this extract (Figure-2). As rotenone is reported to inhibit mitochondrial complex I, trigger ATP depletion, and dopaminergic neuron degeneration; WSRE’s protective effect can be speculated to stem from its positive effect on these very traits. As dopamine as well as WSRE both conferred protection on worms against subsequent rotenone-exposure, WSRE can be thought to have some dopamine-like effect.

**Figure 2.**
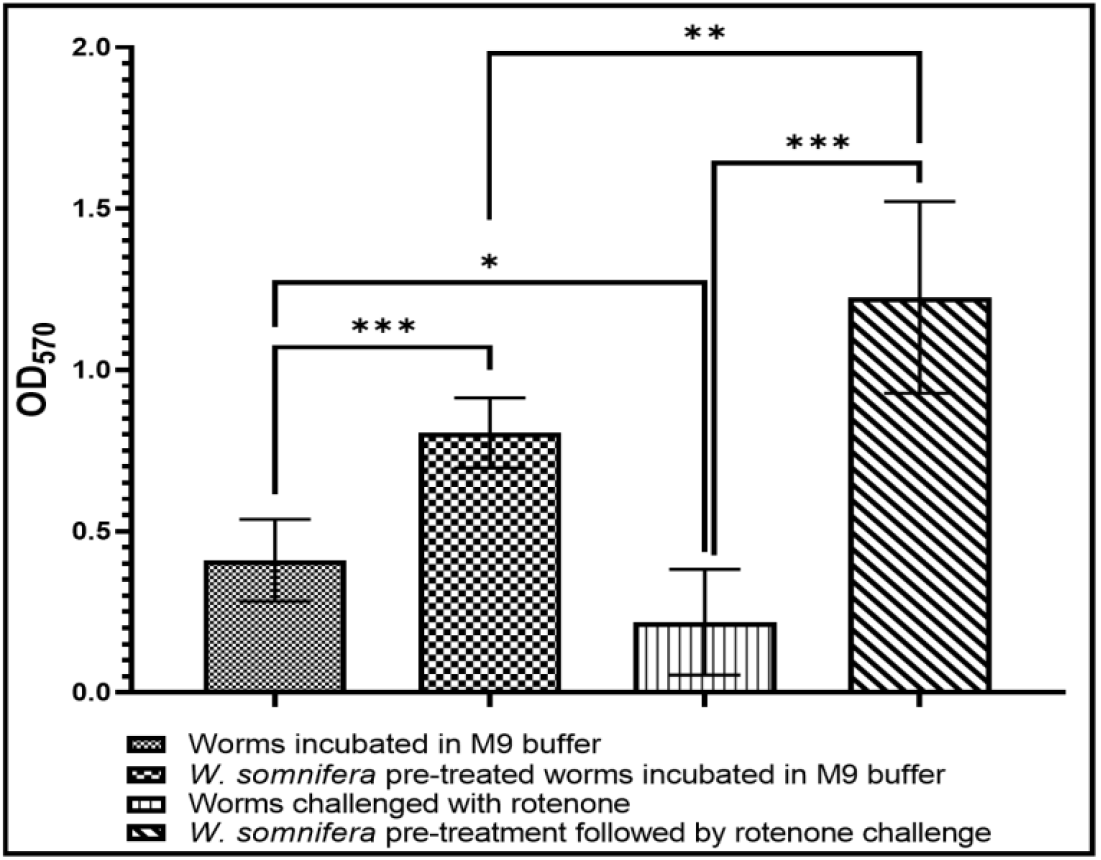
WSRE-pre-fed worms display better metabolic activity in face of rotenone challenge. Alamar Blue^®^ assay was performed on fifth day to capture metabolic activity of worms. The OD_570_ values correspond to the amount of dye reduced by metabolically active worms. WSRE-pre-fed worms displayed higher activity than their counterparts facing or not-facing rotenone challenge. *p ≤ 0.05, **p ≤ 0.01, ***p ≤ 0.001

### 3.2. *W. somnifera* reduces worm’s susceptibility to levamisole

When WSRE- or dopamine-prefed worms were subsequently challenged with levamisole, they were able to maintain a higher worm activity count than their control counterpart with no prior exposure to extract or dopamine. By third day, when levamisole-exposed worms had only 10% activity of that of health control, worms in wells pertaining to WSRE or dopamine exerted 3.77-fold (p=0.0003) and 7.73-fold (p ≤ 0.001) higher activity than control (Figure 3A). Though WSRE pre-feeding reduced worm’s susceptibility to levamisole, it did not support worm activity at par to the health control. Dopamine pre-fed worms could display activity at par to the toxin-non-exposed control worms.

**Figure 3.**
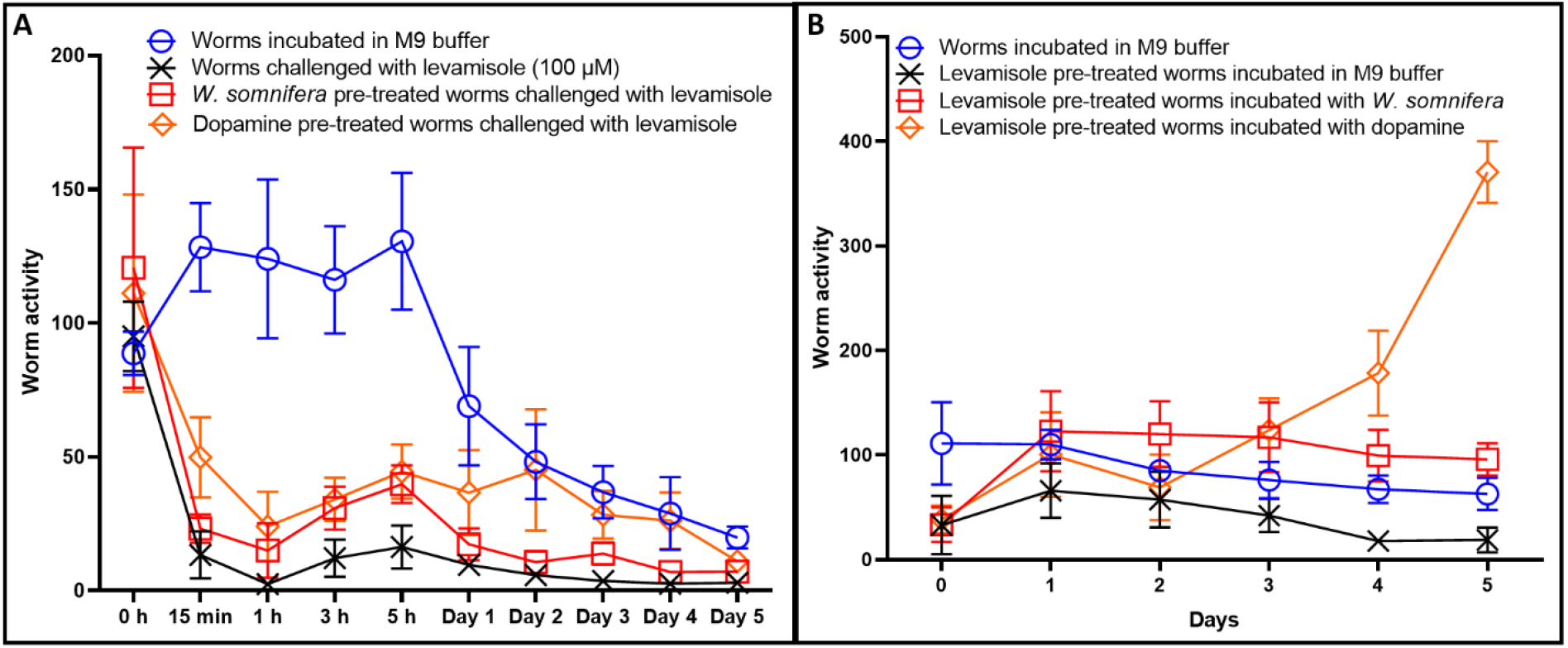
Protective effect of *Withania somnifera* against levamisole. **(A) Prophylactic effect**. Worms were pre-fed for 48 h with WSRE, ascorbic acid, cortisol, or dopamine, followed by continuous exposure to levamisole. Pre-feeding with either dopamine or WSRE noticeably reduced worm susceptibility to levamisole, supporting higher worm activity compared to the levamisole control. See Supplementary Videos D1-D5. This assay was also done with lower levamisole concentrations (25-50 µM), wherein extract-pre-exposure did support faster recovery, however worms were aslo able to recover on their own over a longer period of time (Figure S3A). **(B) Recovery from toxic shock**. Worms were first exposed to levamisole for 5 h, and subsequently allowed to recover in presence of either the WSRE or positive control compounds. WSRE and dopamine, both supported worm recovery from toxic shock, and also supported progeny production on day four onward. See Supplementary Videos E1-E5. For clarity of presentation, lines pertaining to compounds (cortisol or ascorbic acid) showing no activity, which were running parallel to the levamisole control, are not shown in these graphs, so as to reduce visual crowding. WSRE: *Withania somnifera* root extract.

When levamisole-pre-exposed worms were subsequently allowed to recover from the toxic-shock in presence or absence of WSRE, this extract supported recovery faster than dopamine. While WSRE allowed worms to recover and register activity at par to the health control within 24 h, and even surpass (2.09-fold↑; p=0.0003) the health control by 48 h, it took 72 h for dopamine to do so (Figure 3B). At the end of second day, worm activity in WSRE wells was 2.94-fold (p ≤ 0.001) higher than dopamine wells. Toxin-pre-exposed worms in presence of WSRE or dopamine were not only able to recover from the toxic shock, but they could also display fertility, more so in case of dopamine, which contributed to much higher activity counts in dopamine wells on fifth day.

### 3.3. *W. somnifera* reduces worm’s susceptibility to MnCl_**2**_

When worms pre-fed with WSRE were subsequently challenged with MnCl_2_, this toxin was not able to exert any effect on them till first five hours. While control worms lost 65% of activity owing to MnCl_2_ exposure within 15 min, WSRE- or dopamine-pre-fed worms remained unaffected (Figure 4A). Within first three hours of toxin exposure, WSRE-pre-fed worms maintained activity higher than even the health control. While worm activity was almost completely lost in control wells pertaining to MnCl_2_ by end of first day, wells pertaining to WSRE or dopamine pre-exposure still retained measureable activity. By end of fifth day, both, extract- or dopamine-pre-fed worms registered activity at par to those in health control even in face of continuous presence of MnCl_2_.

**Figure 4.**
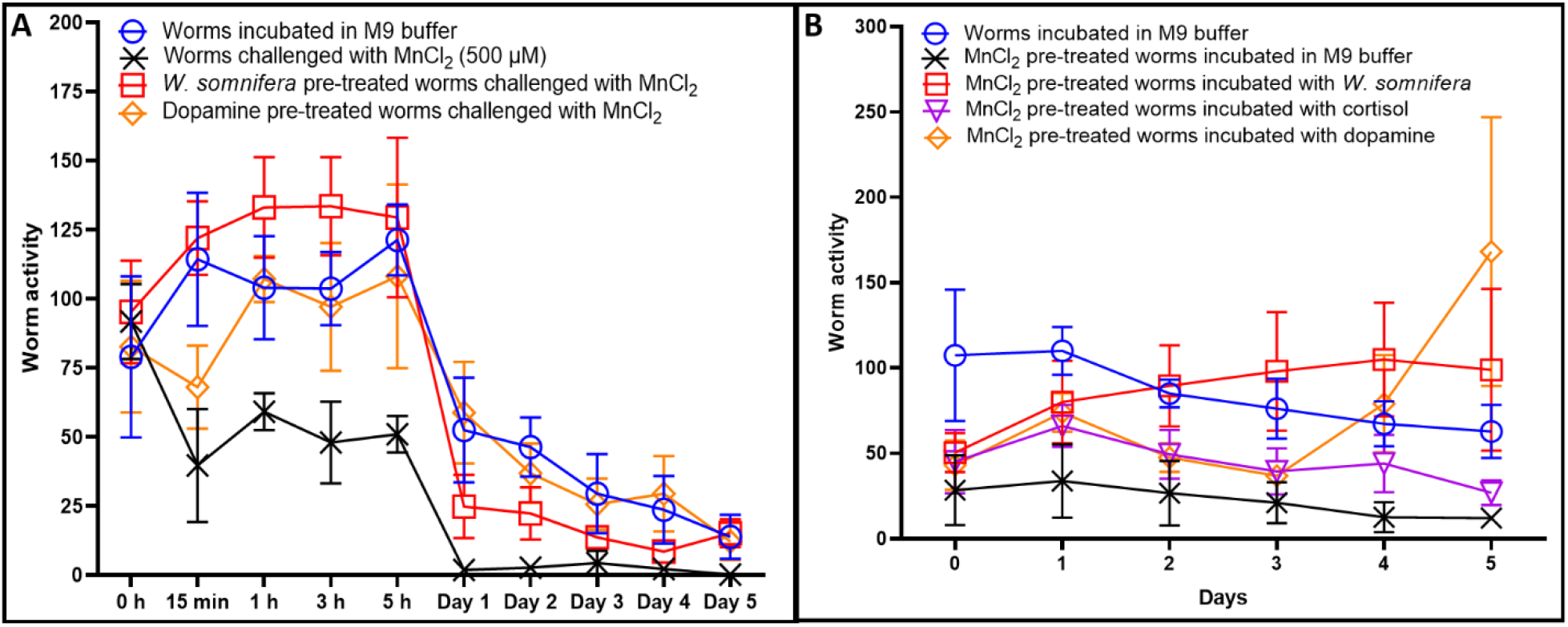
Protective effect of *Withania somnifera* against MnCl_2_. **(A) Prophylactic effect**. Worms were pre-fed for 48 h with WSRE, ascorbic acid, cortisol, or dopamine, followed by continuous exposure to MnCl_2_. Pre-feeding with either dopamine or WSRE noticeably reduced worm susceptibility to MnCl_2_, supporting higher worm activity compared to the MnCl_2_ control. See Supplementary Videos F1-F5. This assay was also done with higher MnCl_2_ concentration (1 mM), wherein extract pre-exposure was able to confer protective effect for lesser duration (Figure S3B). **(B) Recovery from toxic shock**. Worms were first exposed to MnCl_2_ for 3 h, and subsequently allowed to recover in presence of either the WSRE or positive control compounds. WSRE and dopamine, both supported worm recovery from toxic shock. Latter also supported progeny production day four onward. Cortisol also supported moderate recovery, though lower than that observed with WSRE or dopamine. See Supplementary Videos G1-G5. For clarity of presentation, lines pertaining to compounds (cortisol or ascorbic acid) showing no activity, which were running parallel to the MnCl_2_ control, are not shown in these graphs, so as to reduce visual crowding. WSRE: *Withania somnifera* root extract.

When MnCl_2_-pre-exposed worms were subsequently allowed to recover in MnCl_2_-free media supplemented with either WSRE or cortisol or dopamine, their recovery was already started by the end of day one (Figure 4B). Till day three, WSRE supported recovery better than that supported by dopamine. Both dopamine and WSRE restored worm activity to a level higher than the health control. While cortisol also supported worm recovery from MnCl_2_ shock, and its performance was at par to dopamine till day three, fourth day onward dopamine-containing wells registered higher worm activity, which in part stemmed from the progenies.

### 3.4. Worms recover faster from heat shock in presence of WSRE

When wild type worms were challenged with a 3 h heat shock at 37°C in presence or absence of WSRE, and then allowed to recover at 22°C, those in presence of the extract showed faster recovery (Figure 5A). Among different chemicals tried as positive control, only ascorbic acid could support worm recovery from heat shock. While WSRE allowed recovery to start from day two onward, ascorbic acid took three days for the same. By the end of day five, worm activity in wells pertaining to ascorbic acid was at par to that in control wells, while activity in WSRE wells surpassed both of them, partly because progenies appeared in these wells contributing to higher activity counts.

**Figure 5.**
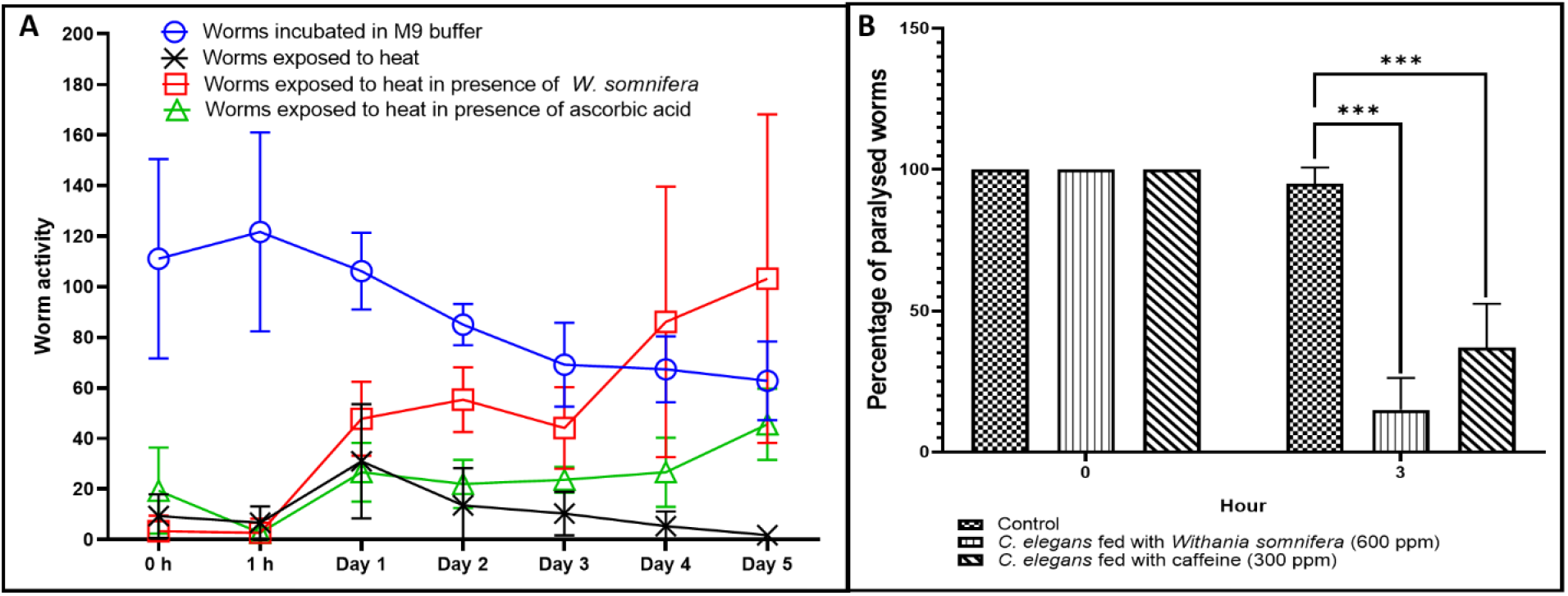
*Withania somnifera* supports *Caenorhabditis elegans* in recovering from heat shock. **(A)** Wild-type worms suspended in liquid media with or without WSRE were first subjected to heat shock (37°C for 3 h), and subsequently allowed to recover at 22°C. Worms incubated in presence of WSRE or ascorbic acid could recover, and displayed higher activity compared to the heat-shocked control worm population. WSRE also allowed the recovering worms to express their fertility. See Supplementary Videos H1-H6. Lines pertaining to dopamine and cortisol are not shown as they did not support worm recovery from heat shock. **(B)** Transgenic worms (CL4176) expressing heat-induced beta-amyloid mediated paralysis were subjected to heat shock (35°C for 4 h) on NGM agar with or without WSRE. Then recovery from heat shock was allowed at 22°C. Worms in presence of WSRE displayed 84±7.07% (p = 0.007) recovery at 3 h end-point post-heat shock. Caffeine was used as positive control, which supported 63.20±15.30% (p=0.01) recovery from paralysis. See Supplementary Videos I1-I5. By 24 h time-point, all worms irrespective of WSRE-exposure were able to recover fully from heat-shock. Results shown in both these graphs were derived from two independent experiments, each with three replicates. WSRE: *Withania somnifera* root extract.

The beta-amyloid expressing transgenic strain CL4176 was also able to exert faster recovery from heat-induced paralysis in presence of WSRE (Figure 5B). Extract’s effect was at par to caffeine (Merck), which was used as positive control (Dostal et al., 2010). This transgenic strain is widely used as a model for neurodegeneration characteristic of Alzheimer’s Disease (Du et al., 2019).

Faster recovery of wild type or transgenic worms from heat shock in presence of WSRE should be viewed in light of the fact that thermosensation and behavioural or locomotory responses to temperature fluctuation in *C. elegans* are not solely physiological in nature, but also a function of neuro-genetic plasticity (Stegeman et al, 2019). It is a neural decision to stop locomotion while facing limits of thermal tolerance, in addition to physiological limits of high temperature stress on locomotory activity. Since the thermal performance of *C. elegans* is sensitive to genetic perturbations of behavioural decision-making as well as physiological limits, WSRE’s protective activity in face of thermal challenge can be considered a clue to its wide spectrum of beneficial activities including neuroprotective.

This study has found WSRE to offer protection to worms challenged with three different toxins―rotenone, MnCl_2_, and levamisole. Though we did not perform any mechanistic assays, based on known modes of action of these toxins, it may be speculated that WSRE has neuroprotective as well as mitoprotective activity. Rotenone (Subaraja and Janardhanam, 2019) and MnCl_2_ (Chen et al., 2015) are known neurotoxins, which can trigger mitochondrial dysfunction as well as degeneration of dopaminergic neurons (Huang et al., 2021; Ibarra-Gutiérrez et al., 2023), conditions which are considered among hallmarks of neurodegenerative diseases like Parkinson’s disease (Pfeifer, 2024).

Levamisole induces neuromuscular paralysis by interfering with acetylcholine neurotransmission (Lewis et al., 1980). Since levamisole is believed to act on multiple receptors, with multiple genes being responsible for coding receptor ion-channels, and the target sites are pharmacologically diverse (Martin et al., 2005), WSRE can be logically believed to be offering protection against this toxin by displaying a multiplicity of targets in worms, which is expected from any natural product containg multiple active ingredients.

WSRE’s protective effect against MnCl_2_-induced toxicity becomes more relevant in light of the fact that environmental exposure to manganese is among the risk factors for the occurrence of Parkinson’s disease. Manganism, a disease with characteristic degeneration of dopamine neurons, has clinical manifestations similar to Parkinson’s disease. Dopamine-like effect of WSRE on MnCl_2_-challenged worms can partly be explained by the observation that dopamine influences the sensitivity to manganese-induced neurotoxicity in *C. elegans* (Raj and Thekkuveettil, 2024).

Results of the present study supporting the claims of adaptogenic/anti-stress potential of WSRE corroborate well with earlier such studies conducted in different model organisms or with human volunteers. For example, Salve et al. (2019) reported adaptogenic and anxiolytic effects of ashwagandha root extract in healthy adults at 250-600 mg/day. Lopresti et al. (2019) reported stress-relieving effect of WSRE at 240 mg/day in human adults. Manjunath and Muralidhara (2015) found WSRE to markedly offset rotenone-induced locomotor deficits, oxidative impairments and neurotoxicity in *Drosophila melanogaster*.

As WSRE-pre-exposure enhanced worm’s capacity to withstand subsequent constant presence of these toxins, as well as, their capacity to recover from toxic shock induced by either of these toxins, and heat-shock too, WSRE’s anti-stress spectrum can be said to be fairly wide (Figure 6). Against heat stress, WSRE’s protective action was similar to ascorbic acid, while against chemical stressors, this extract’s effect appeared to be dopamine-like. Further, WSRE also displayed protective potential against beta-amyloid peptide toxicity in transgenic worm. It should also be noted that in our experimental set-up, the worms used were gnotobiotic and were not provided any bacterial food during assay to avoid any possible confounding effect of worm microbiota on results. That means the worms were under an additional stress of restricted nutrient availability. WSRE’s beneficial effect should be appreciated in context of this dual stress i.e. stressor-challenge in oligotrophic conditions. This extract was found to retain its positive effect on worm healthspan even after refrigerated storage of thirteen months (Figure S4). Its multimode protective potential is not surprising given it being a natural product with multiple ingredients, quite a few of whom acting as active principles. Our results support the nutraceutical candidature of *W. somnifera* root extracts, and the need for further investigation into the molecular mechanisms associated with its biological activities.

**Figure 6.**
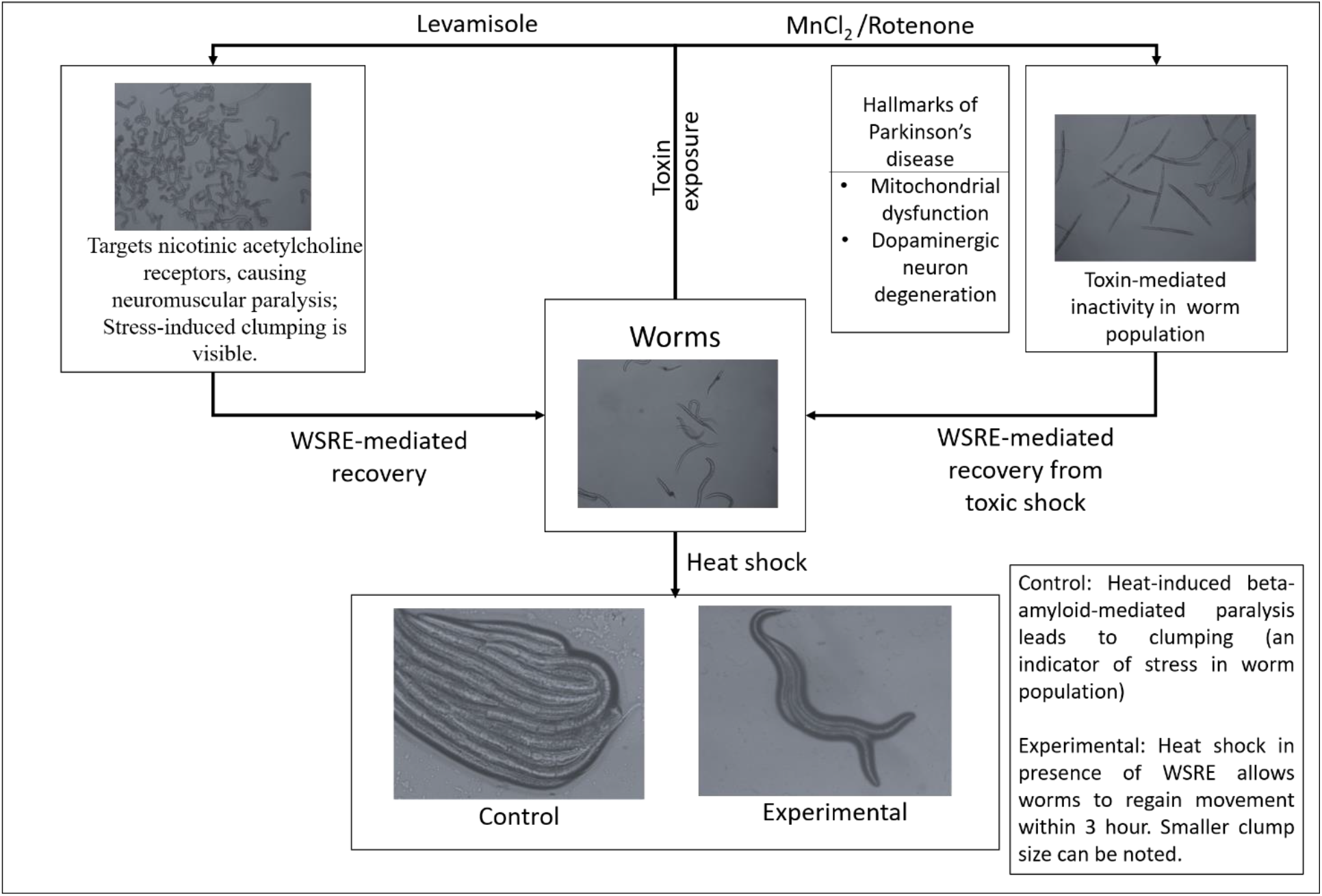
Schematic summary of adaptogenic effect of *W. somnifera* root extract.

## Supplementary files

Table S1-S6

Figure S1-S4

Videos A1-I5 (legend to these videos provided as Appendix-A)

## Acknowledgement

Authors thank NERF (Nirma Education and Research Foundation), Ahmedabad, for infrastructural and financial support, and for providing a doctoral stipend to NT. NT acknowledges support from the Government of Gujarat via their SHODH scheme. N2 Bristol strain was provided by the CGC, which is funded by NIH Office of Research Infrastructure Programs (P40 OD010440). Gemini Gajera is thanked for assistance with optimization of toxin dosage. Xavier’s Research Foundation, Ahmedabad is thanked for providing the transgenic worm strain.

## References

1. Johri A, Beal MF. Mitochondrial dysfunction in neurodegenerative diseases. The Journal of Pharmacology and Experimental Therapeutics. 2012;342(3):619–30. 10.1124/jpet.112.192138

2. Vaidya VG, Naik NN, Ganu G, Parmar V, Jagtap S, Saste G, Bhatt A, Mulay V, Girme A, Modi SJ, Hingorani L. Clinical pharmacokinetic evaluation of Withania somnifera (L.) Dunal root extract in healthy human volunteers: a non-randomized, single dose study utilizing UHPLC-MS/MS analysis. Journal of Ethnopharmacology. 2024; 322:117603. 10.1016/j.jep.2023.117603

3. Saha P, Ajgaonkar S, Maniar D, Sahare S, Mehta D, Nair S. Current insights into transcriptional role (s) for the nutraceutical Withania somnifera in inflammation and aging. Frontiers in Nutrition. 2024; 11:1370951. 10.3389/fnut.2024.1370951

4. Verpoorte R. New times for traditional medicine research. Journal of Ethnopharmacology. 2017;100(197):1. 10.1016/j.jep.2017.01.018

5. Thakkar N, Gajera G, Mehta D, Nair S, Kothari V. Withania somnifera root extract (LongeFera™) confers beneficial effects on health and lifespan of the model worm Caenorhabditis elegans. Drug Target Insights. 2025; 19:18. doi: 10.33393/dti.2025.3368

6. Deji-Oloruntoba OO, Elufioye TO, Adefegha SA, Jang M. Can Caenorhabditis elegans serve as a reliable model for drug and nutraceutical discovery? Applied Biosciences. 2025;4(2):23. 10.3390/applbiosci4020023

7. Sammi SR, Jameson LE, Conrow KD, Leung MC, Cannon JR. Caenorhabditis elegans neurotoxicity testing: novel applications in the adverse outcome pathway framework. Frontiers in Toxicology. 2022; 4:826488. doi: 10.3389/ftox.2022.826488

8. Xu H, Wang Q, Zhou Y, Chen H, Tao J, Huang J, Miao Y, Zhao J, Wang Y. Cremastra appendiculata Polysaccharides Alleviate Neurodegenerative Diseases in Caenorhabditis elegans: Targeting Amyloid-β Toxicity, Tau Toxicity and Oxidative Stress. International Journal of Molecular Sciences. 2025;26(8):3900. 10.3390/ijms26083900

9. Corsi AK, Wightman B, Chalfie M. A transparent window into biology: a primer on Caenorhabditis elegans. Genetics. 2015;200(2):387–407. 10.1534/genetics.115.180133

10. Gonzalez-Hunt CP, Luz AL, Ryde IT, Turner EA, Ilkayeva OR, Bhatt DP, Hirschey MD, Meyer JN. Multiple metabolic changes mediate the response of Caenorhabditis elegans to the complex I inhibitor rotenone. Toxicology. 2021; 447:152630. 10.1016/j.tox.2020.152630

11. Bora S, Vardhan GS, Deka N, Khataniar L, Gogoi D, Baruah A. Paraquat exposure over generation affects lifespan and reproduction through mitochondrial disruption in C. elegans. Toxicology. 2021; 447:152632. 10.1016/j.tox.2020.152632

12. Page KE, White KN, McCrohan CR, Killilea DW, Lithgow GJ. Aluminium exposure disrupts elemental homeostasis in Caenorhabditis elegans. Metallomics. 2012;4(5):512-22. DOI: 10.1039/c2mt00146b

13. Moline B. Characterization of manganese-induced neurodegenration in C. elegans treated with winterberry leaf extract.

14. Chen P, Chakraborty S, Peres TV, Bowman AB, Aschner M. Manganese-induced neurotoxicity: from C. elegans to humans. Toxicology Research. 2015;4(2):191–202. 10.1039/c4tx00127c

15. Hernando G, Bouzat C. Caenorhabditis elegans neuromuscular junction: GABA receptors and ivermectin action. PLoS One. 2014;9(4):e95072. 10.1371/journal.pone.0095072

16. Martin RJ, Robertson AP, Buxton SK, Beech RN, Charvet CL, Neveu C. Levamisole receptors: a second awakening. Trends in Parasitology. 2012;28(7):289–96.

17. Patil VM, Verma S, Masand N. Prospective mode of action of Ivermectin: SARS-CoV-2. European Journal of Medicinal Chemistry Reports. 2022; 4:100018. 10.1016/j.ejmcr.2021.100018

18. Holden-Dye L, Walker R. Anthelmintic drugs and nematocides: studies in Caenorhabditis elegans. WormBook: the online review of C. elegans biology. 2014:1–29. 10.1895/wormbook.1.143.2

19. Fu X, Tang Y, Dickinson BC, Chang CJ, Chang Z. An oxidative fluctuation hypothesis of aging generated by imaging H_2_O_2_ levels in live Caenorhabditis elegans with altered lifespans. Biochemical and Biophysical Research Communications. 2015;458(4):896–900. 10.1016/j.bbrc.2015.02.055

20. Yasuda K, Kubo Y, Murata H, Sakamoto K. Cortisol promotes stress tolerance via DAF-16 in Caenorhabditis elegans. Biochemistry and Biophysics Reports. 2021; 26:100961. 10.1016/j.bbrep.2021.100961

21. Chou SH, Chen YJ, Liao CP, Pan CL. A role for dopamine in C. elegans avoidance behavior induced by mitochondrial stress. Neuroscience Research. 2022; 178:87–92. 10.1016/j.neures.2022.01.005

22. Harrington LA, Harley CB. Effect of vitamin E on lifespan and reproduction in Caenorhabditis elegans. Mechanisms of Ageing and Development. 1988;43(1):71–8. 10.1016/0047-6374(88)90098-X

23. Dostal V, Roberts CM, Link CD. Genetic mechanisms of coffee extract protection in a Caenorhabditis elegans model of β-amyloid peptide toxicity. Genetics. 2010;186(3):857–66. 10.3390/molecules24040729

24. Springer JE, Azbill RD, Carlson SL. A rapid and sensitive assay for measuring mitochondrial metabolic activity in isolated neural tissue. Brain Research Protocols. 1998;2(4):259–63. 10.1016/S1385-299X(97)00045-7

25. Du F, Zhou L, Jiao Y, Bai S, Wang L, Ma J, Fu X. Ingredients in Zijuan Pu’er tea extract alleviate β-amyloid peptide toxicity in a Caenorhabditis elegans model of Alzheimer’s disease likely through DAF-16. Molecules. 2019;24(4):729. 10.1534/genetics.110.120436

26. Stegeman GW, Medina D, Cutter AD, Ryu WS. Neuro-genetic plasticity of Caenorhabditis elegans behavioral thermal tolerance. BMC Neuroscience. 2019;20(1):26. 10.1186/s12868-019-0510-z

27. Subaraja M, Janardhanam Vanisree A. Aberrant neurotransmissional mRNAs in cerebral ganglions of rotenone-exposed Lumbricus terrestris exhibiting motor dysfunction and altered cognitive behavior. Environmental Science and Pollution Research. 2019;26(14):14461–72. 10.1007/s11356-019-04740-y

28. Huang Y, Wen Q, Huang J, Luo M, Xiao Y, Mo R, Wang J. Manganese (II) chloride leads to dopaminergic neurotoxicity by promoting mitophagy through BNIP3-mediated oxidative stress in SH-SY5Y cells. Cellular & Molecular Biology Letters. 2021;26(1):23. 10.1186/s11658-021-00267-8

29. Ibarra-Gutiérrez MT, Serrano-García N, Orozco-Ibarra M. Rotenone-induced model of Parkinson’s disease: Beyond mitochondrial complex I inhibition. Molecular Neurobiology. 2023 Apr;60(4):1929–48. 10.1007/s12035-022-03193-8

30. Pfeifer GP. DNA damage and Parkinson’s disease. International Journal of Molecular Sciences. 2024;25(8):4187. 10.3390/ijms25084187

31. Lewis JA, Wu CH, Levine JH, Berg H. Levamisole-resitant mutants of the nematode Caenorhabditis elegans appear to lack pharmacological acetylcholine receptors. Neuroscience. 1980;5(6):967–89. 10.1016/0306-4522(80)90180-3

32. Martin RJ, Verma S, Levandoski M, Clark CL, Qian H, Stewart M, Robertson AP. Drug resistance and neurotransmitter receptors of nematodes: recent studies on the mode of action of levamisole. Parasitology. 2005;131(S1): S71–84. doi:10.1017/S0031182005008668

33. Raj V, Thekkuveettil A. Dopamine controls the Sensitivity to Manganese induced Dopaminergic neurotoxicity in Caenorhabditis elegans. bioRxiv. 2024:2024-10. 10.1101/2024.10.23.619952

34. Salve J, Pate S, Debnath K, Langade D, Langade DG. Adaptogenic and anxiolytic effects of ashwagandha root extract in healthy adults: a double-blind, randomized, placebo-controlled clinical study. Cureus.;11(12). DOI 10.7759/cureus.6466

35. Lopresti AL, Smith SJ, Malvi H, Kodgule R. An investigation into the stress-relieving and pharmacological actions of an ashwagandha (Withania somnifera) extract: A randomized, double-blind, placebo-controlled study. Medicine. 2019;98(37):e17186. 10.1097/MD.0000000000017186

36. Manjunath MJ, Muralidhara. Standardized extract of Withania somnifera (Ashwagandha) markedly offsets rotenone-induced locomotor deficits, oxidative impairments and neurotoxicity in Drosophila melanogaster. Journal of Food Science and Technology. 2015;52(4):1971–81. DOI 10.1007/s13197-013-1219-0

