## Supplementary Tables, Figures, and Videos for "*Withania somnifera* root extract reduces susceptibility of the model worm *Caenorhabditis elegans* to different types of stressors": Appendix A_Legend to videos.pdf

### Legend to Supplementary Videos

for

Videos were captured using a Magnus Camera (5.1 MP) attached to a binocular microscope (Labomed Vision 2000) equipped with halogen light source. While all videos were captured observing through 4X objective, videos I1-I5 were captured using 10X objective.

**Assays with chemical stressors**

**A1-A2**: Health control for assays pertaining to the prophylactic potential of WSRE. Gnotobiotic worm in M9 buffer (devoid of plant extract or toxin) captured at day zero (A1) and day 4 (A2). Active movement is visible.

**A3-A4:** Health control for assays pertaining to therapeutic potential of WSRE, post-toxin exposure. Worms captured on day one (A3). Worms captured on day three (A4).

**B1**: Worms challenged with rotenone (3 µM) captured after 60 min of toxic shock. **B2**: Extract (600 ppm)-pre-exposed worms challenged with rotenone captured after 60 min of toxic shock. **B3**: Dopamine-pre-exposed worms challenged with rotenone captured after 60 min of toxin shock. Dopamine (5mM) pre-exposure seems to have a positive effect on worm morphology too.

**C1**: Worms pre-exposed (5 h) to rotenone captured at day two. Most of them have lost activity. **C2:** Rotenone-pre-exposed worms allowed to recover in presence of WSRE captured on day two. **C3**: Rotenone-pre-exposed worms captured at day five. **C4:** Rotenone-pre-exposed worms allowed to recover in presence of WSRE captured on day five. Progeny is also visible. **C5**: Rotenone-pre-exposed worms allowed to recover in presence of dopamine, captured on day five. Vigorous movement, bigger morphology and progeny can be visualized.

**D1**: Worms challenged with levamisole (100 µM), captured at 5 h post-toxin challenge. **D2:** Extract-pre-exposed worms challenged with levamisole, captured at 5 h post-toxin challenge. **D3**: Worms challenged with levamisole, captured on day-four. Slow movement, and clumping (indication of them being under stress) is visible. **D4**: Extract-pre-fed worms challenged with levamisole, captured on day-four. Clumping is absent, as these worms are able to travel farther from each-other owing to prophylactic protection offered by WSRE, and are under lesser stress. **D5:** Dopamine-pre-fed worms challenged with levamisole, captured on day-four. Degree of clumping is lesser than toxicity control (D3), but more than WSRE-pre-fed worms (D4).

**E1:** Worms pre-exposed to levamisole, captured on day one. **E2:** Levamisole pre-exposed worms subsequently allowed to recover in presence of WSRE, captured on day one. **E3**: Worms pre-exposed to levamisole, captured on day five. Clumping owing to reduced ability to move farther distances is visible. **E4**: Worms pre-exposed to levamisole, allowed to recover in presence of WSRE, captured on day five. No clumping of worms is observed, as they are able to move farther distances. Progeny worms are also visible. **E5**: Worms pre-exposed to levamisole, allowed to recover in presence of dopamine, captured on day five. Vigorous movement, and many progenies are visible.

**F1**: Worms challenged with $MnCl_2$ (500 µM) captured 5 h post-toxin challenge. **F2**: Extract-pre-fed worms challenged with $MnCl_2$ captured 5 h post-toxin challenge. **F3:** Worms challenged with $MnCl_2$ captured on day two, have lost all the activity. **F4**: Extract-pre-fed worms challenged with $MnCl_{2,}$ captured on day two. Few live worms are visible. **F5:** Dopamine-pre-fed worms challenged with $MnCl_{2,}$ captured on day two.

**G1:** $MnCl_2$-pre-exposed (for 3 hour) worms, captured on day three. **G2:** $MnCl_2$-pre-exposed worms allowed to recover in presence of WSRE, captured on day three. Activity is better than health control (**A4**). **G3**: $MnCl_2$-pre-exposed worms captured on day five. **G4**: $MnCl_2$-pre-exposed worms allowed to recover in presence of WSRE, captured on day five. **G5**: $MnCl_2$-pre-exposed worms allowed to recover in presence of dopamine, captured on day five. Vigorous movement, bigger morphology, and progenies can be observed.

**Thermorecovery assays**

**H1**: Wild type worms in liquid media captured immediately after a 3 h heat shock at 37°C. Those subject to heat shock in presence of WSRE, not shown here, were also similarly devoid of any movement.; Wild type worms incubated at 22°C in absence **(H2)** or presence **(H3)** of WSRE, allowed to recover from previous heat shock, captured on day two. Better movement in H3 than H2 is visible. Corresponding video for vitamin C is **H4**; On day five, while almost all heat-pre-exposed control worms were dead **(H5),** active movement in wells corresponding to WSRE **(H6)** or vitamin-C **(H7)** was visible. Progenies are also visible in **H6**.

Transgenic worms on NGM agar expressing heat-induced paralytic phenotype captured immediately after 4 h heat shock at 35°C, in absence **(I1)** or presence of WSRE **(I2).** Slow movement and stress-induced clumping can be observed in both cases; When worms were allowed to recover from heat shock by incubating at 22°C, at three-hour time point observation, their recovery was better than control **(I3)** in presence of WSRE **(I4)** or caffeine **(I5).** Reduced clumping of worms in I4-I5 is indication of lesser stress.
