## Supplementary Tables, Figures, and Videos for "*Withania somnifera* root extract reduces susceptibility of the model worm *Caenorhabditis elegans* to different types of stressors": Supplementary Tables and Figures_C. elegans_W. somnifera_Anti-stress_Dec 2025_VK.pdf

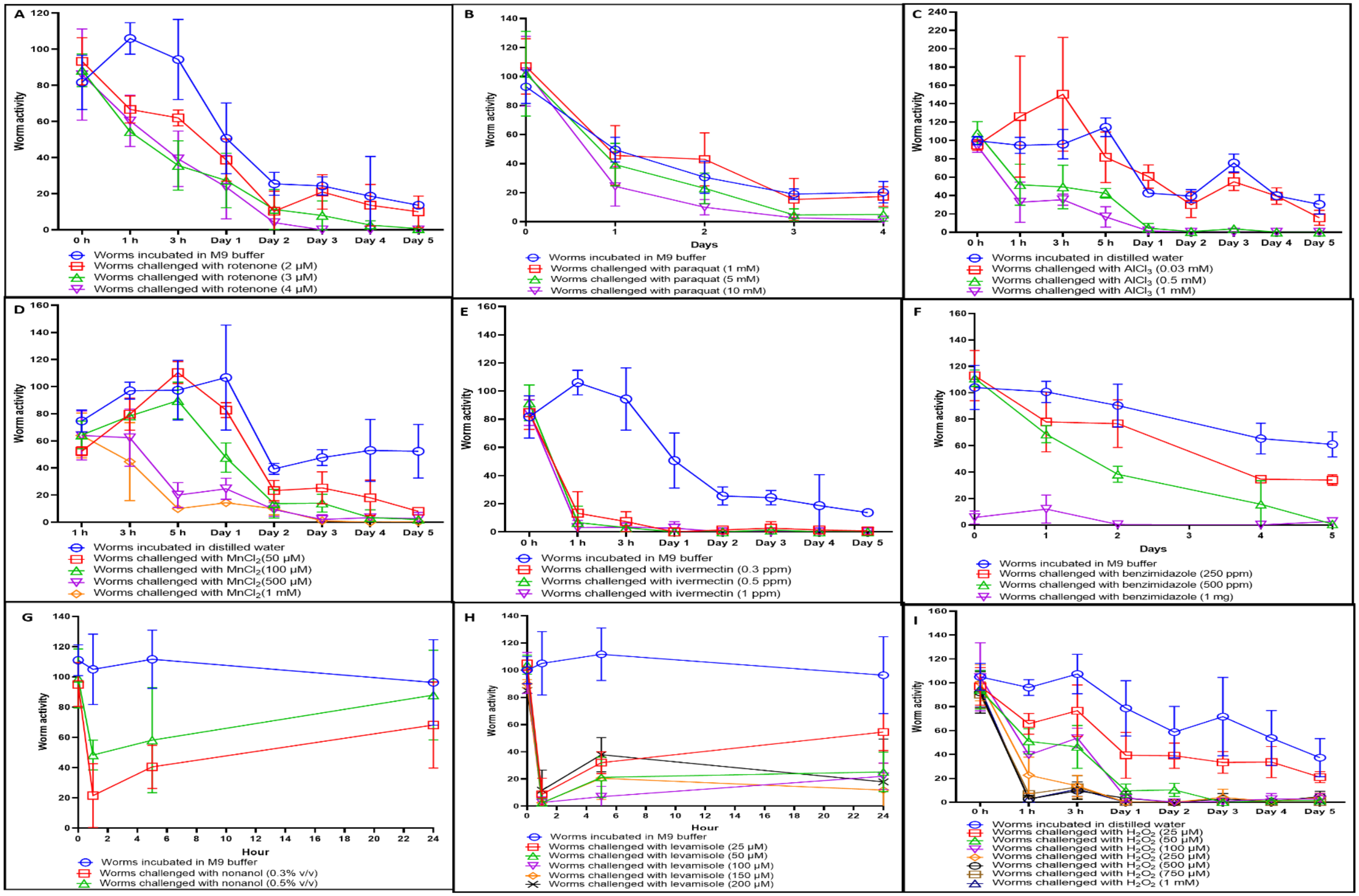


**Figure S1. Effect of toxic chemicals on *C. elegans*** **(A)** Rotenone **(B)** Paraquat **(C)** Aluminum Chloride ($AlCl_3$) **(D)** Manganese chloride ($MnCl_2$) **(E)** Ivermectin **(F)** Benzimidazole **(G)** Nonanol **(H)** Levamisole **(I)** Hydrogen peroxide ($H_2O_2$). Population size of worms per well was approximately 100.

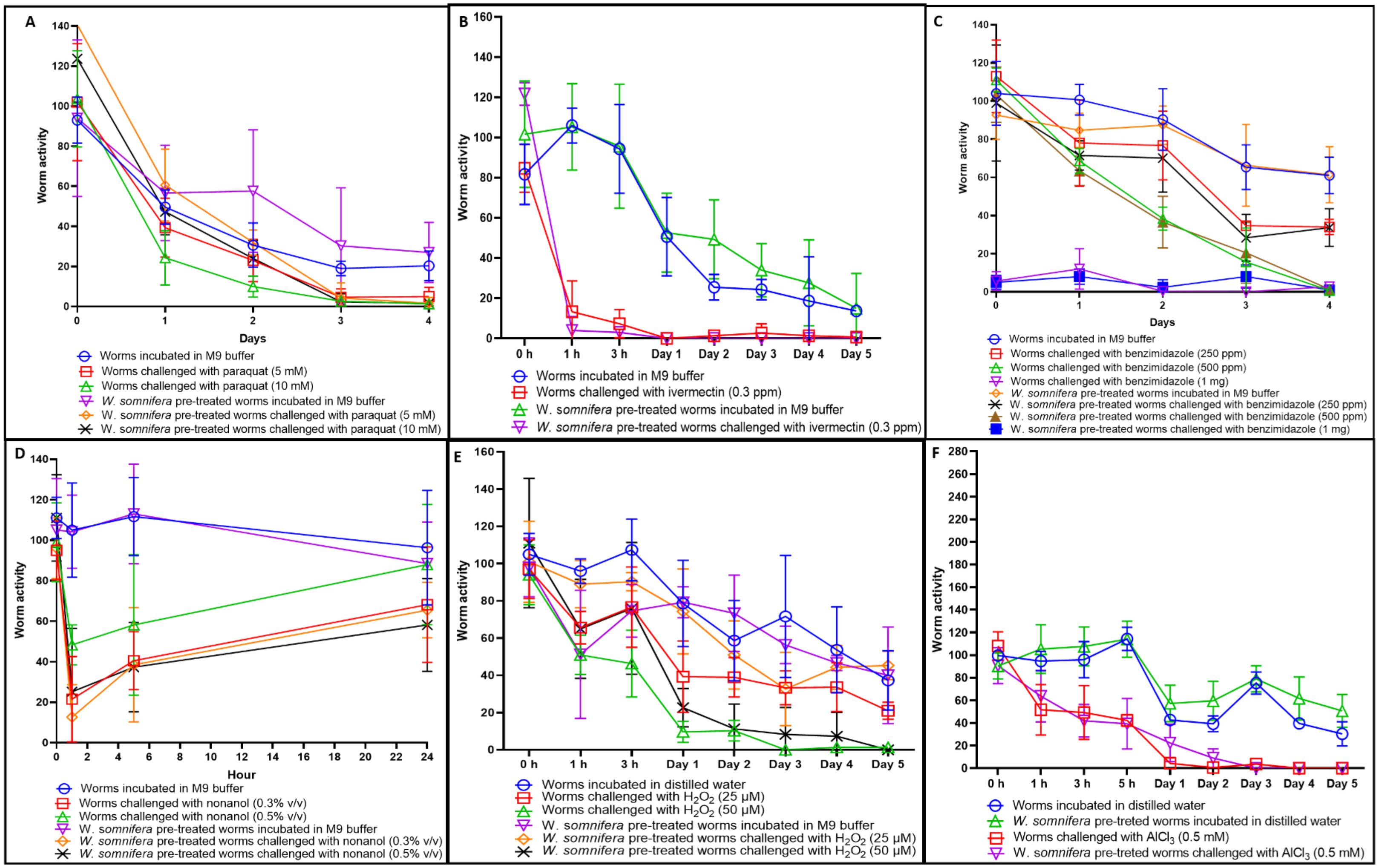


**Figure S2. Pre-treatment with *Withania somnifera* did not protect worms from subsequent challenge with (A)** Paraquat **(B)** Ivermectin **(C)** Benzimidazole **(D)** Nonanol **(E)** Hydrogen peroxide ($H_2O_2$) **or (F)** Aluminum Chloride ($AlCl_3$)

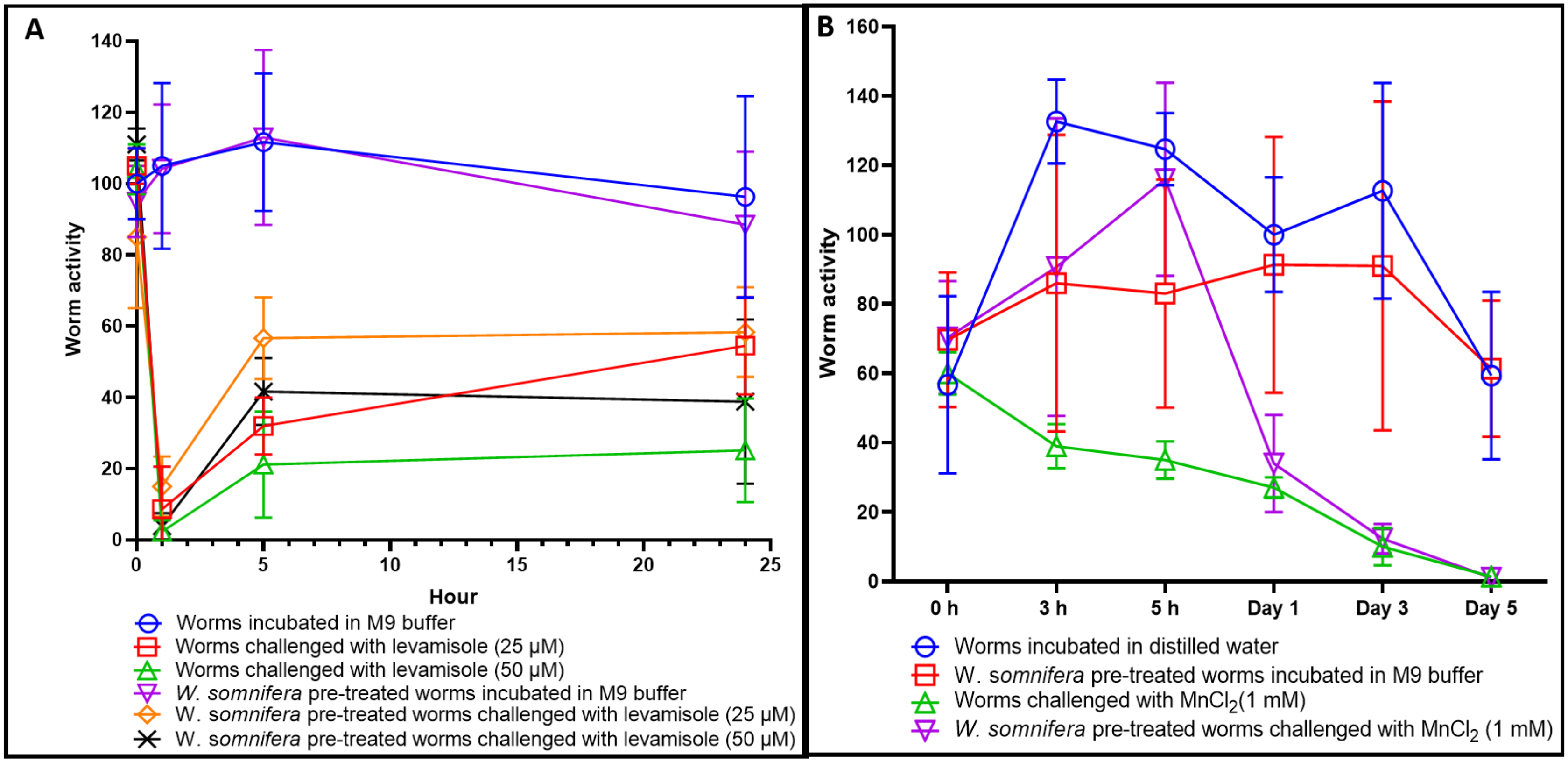


**Figure S3. *Withania somnifera* offers prophylactic protection to worms subsequently challenged with stressors. (A) Levamisole.** Worms were pre-fed for 48 h with WSRE, followed by continuous exposure to levamisole. Pre-feeding with WSRE supported higher worm activity compared to the levamisole control till 5 hour. After that worm activity was statistically equal in all wells (data not shown). **(B) Manganese chloride ($MnCl_2$).** Worms were pre-fed for 48 h with WSRE, followed by continuous exposure to $MnCl_2$. Pre-feeding with WSRE delayed the death in worm population.

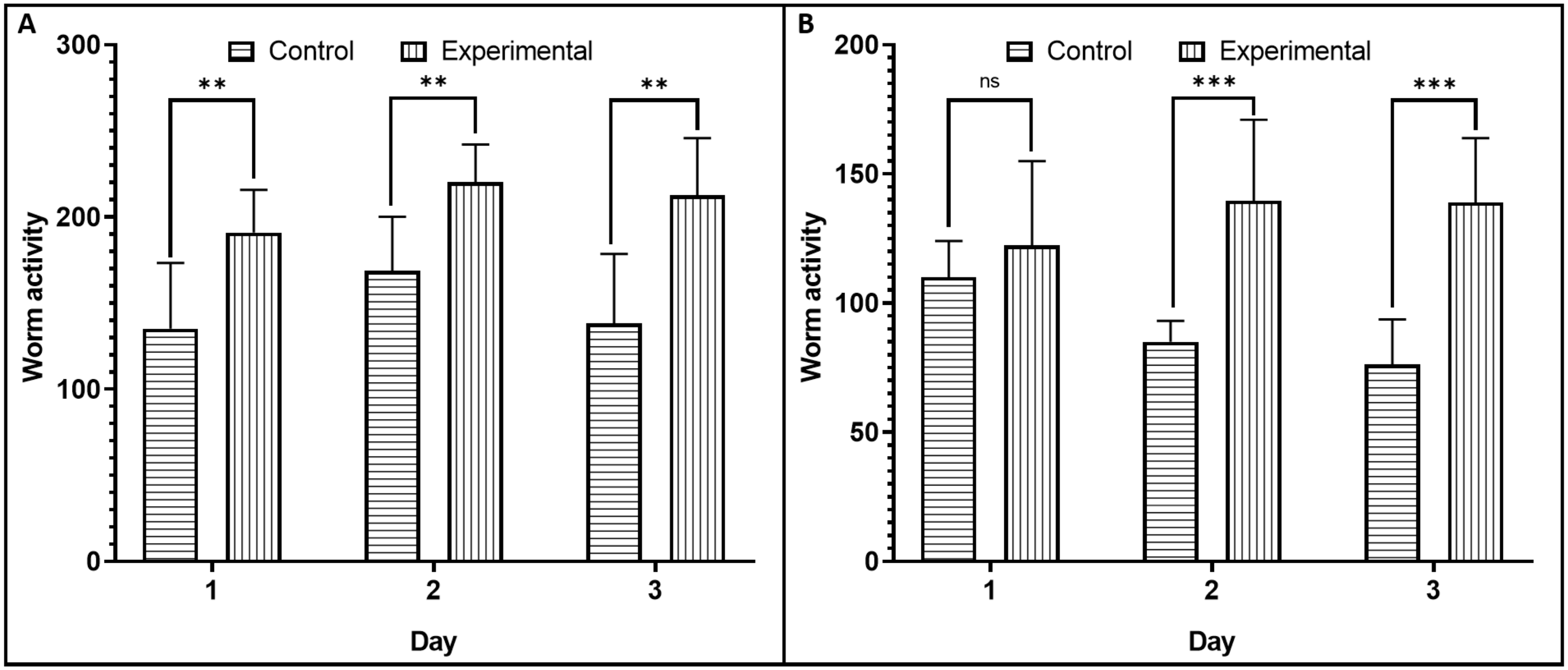


**Figure S4. *Withania somnifera* extract retains its beneficial effect on worm healthspan even after one year of refrigerated storage. (A)** Study conducted in July 2024. **(B)** Study conducted in August 2025**.** WSRE dissolved in water, filter-sterilized, and stored under refrigeration in glass vial**.** Population size of worms was 100 per well. **$p \leq 0.01$, ***$p \leq 0.001$

Table S1-S6 contains the mean values of absolute worm activity counts as measured by the worm tracker, from which the figures provided in the main manuscript were generated.

### Table S1. Protective effect of *Withania somnifera* against rotenone [Raw data pertaining to Figure 1(A)]

| | | 0 h | 15 min | 1 h | 3 h | 5 h | Day 1 | Day 2 | Day 3 | Day 4 | Day 5 |
|---|---|---|---|---|---|---|---|---|---|---|---|
| A | Worms incubated in M9 buffer | 88.67 ± 8.14 | 128.33 ± 16.46 | 124.00 ± 29.64 | 116.17 ± 20.06 | 130.50 ± 25.62 | 69.00 ± 22.14 | 48.17 ± 13.96 | 36.83 ± 9.81 | 28.83 ± 13.59 | 19.83 ± 4.07 |
| B | Worms challenged with rotenone | 88.83 ± 11.65 | 47.00 ± 16.25[a] | 44.17 ± 10.15[a] | 42.67 ± 12.23[a] | 38.50 ± 10.09[a] | 15.17 ± 5.19[a] | 5.17 ± 5.74[a] | 3.50 ± 4.18[a] | 2.67 ± 4.13[a] | 0.00 ± 0.00[a] |
| C | *W. somnifera* pre-treated worms challenged with rotenone | 99.33 ± 13.60 | 96.00 ± 21.14[b] | 101.33 ± 13.75[b] | 70.83 ± 8.28[b] | 61.17 ± 11.36[b] | 39.17 ± 5.85[b] | 24.67 ± 11.50[b] | 12.50 ± 1.76[b] | 10.67 ± 3.93[b] | 5.33 ± 1.86[b] |
| D | Dopamine pre-treated worms challenged with rotenone | 90.33 ± 20.88 | 120.67 ± 14.14[c] | 116.00 ± 30.71[c] | 108.50 ± 14.52[c] | 92.50 ± 31.87[c] | 57.00 ± 6.07[c] | 37.67 ± 9.29[c] | 21.83 ± 3.54[c] | 20.83 ± 5.56[c] | 12.67 ± 3.20[c] |

Values presented are Mean ± SEM. [a]: Denotes statistically significant difference between A-B; [b]: Denotes statistically significant difference between B-C. [c]: Denotes statistically significant difference between B-D.

### Table S2. Protective effect of *Withania somnifera* against rotenone [Raw data pertaining to Figure 1(B)]

| | | 0 h | Day 1 | Day 2 | Day 3 | Day 4 | Day 5 |
|---|---|---|---|---|---|---|---|
| A | Worms incubated in M9 buffer | 111.11 ± 39.51 | 110.00 ± 14.10 | 85.00 ± 8.14 | 76.22 ± 17.51 | 67.33 ± 13.09 | 62.78 ± 15.54 |
| B | Rotenone pre-treated worms incubated in M9 buffer | 37.56 ± 20.11[a] | 64.67 ± 18.29[a] | 49.78 ± 11.63[a] | 46.00 ± 15.09[a] | 31.44 ± 12.12[a] | 24.33 ± 12.47[a] |
| C | Rotenone pre-treated worms incubated with *W. somnifera* | 55.89 ± 34.45 | 132.44 ± 51.64[b] | 145.33 ± 54.64[b] | 133.89 ± 45.13[b] | 119.78 ± 47.25[b] | 135.11 ± 46.47[b] |
| D | Rotenone pre-treated worms incubated with cortisol | 34.22 ± 13.27 | 90.00 ± 30.96[c] | 65.78 ± 12.87[c] | 62.11 ± 11.13[c] | 52.33 ± 17.16[c] | 47.00 ± 11.16[c] |
| E | Rotenone pre-treated worms incubated with dopamine | 36.00 ± 16.82 | 105.00 ± 20.78[d] | 75.44 ± 9.06[d] | 108.22 ± 16.20[d] | 126.22 ± 30.31[d] | 201.89 ± 78.50[d] |

Values presented are Mean ± SEM. [a]: Denotes statistically significant difference between A-B; [b]: Denotes statistically significant difference between B-C; [c]: Denotes statistically significant difference between B-D; [d]: Denotes statistically significant difference between B-E.

### Table S3. Protective effect of *Withania somnifera* against levamisole [Raw data pertaining to Figure 3(A)]

| | | 0 h | 15 min | 1 h | 3 h | 5 h | Day 1 | Day 2 | Day 3 | Day 4 | Day 5 |
|---|---|---|---|---|---|---|---|---|---|---|---|
| A | Worms incubated in M9 buffer | 88.67 ± 8.14 | 128.33 ± 16.46 | 124.00 ± 29.64 | 116.17 ± 20.06 | 130.50 ± 25.62 | 69.00 ± 22.14 | 48.17 ± 13.96 | 36.83 ± 9.81 | 28.83 ± 13.59 | 19.83 ± 4.07 |
| B | Worms challenged with levamisole | 95.00 ± 12.98 | 13.33 ± 8.76[a] | 2.50 ± 3.89[a] | 12.17 ± 6.91[a] | 16.33 ± 8.02[a] | 9.67 ± 2.58[a] | 5.83 ± 3.71[a] | 3.67 ± 2.16[a] | 2.67 ± 2.16[a] | 3.00 ± 2.83[a] |
| C | *W. somnifera* pre-treated worms challenged with levamisole | 120.67 ± 44.88 | 23.17 ± 5.31[b] | 15.00 ± 10.20[b] | 30.83 ± 8.06[b] | 39.83 ± 7.08[b] | 17.33 ± 5.92[b] | 10.67 ± 4.08[b] | 13.83 ± 4.17[b] | 7.00 ± 3.03[b] | 7.17 ± 1.72[b] |
| D | Dopamine pre-treated worms challenged with levamisole | 111.17 ± 36.88 | 49.83 ± 14.97[c] | 24.00 ± 12.96[c] | 34.17 ± 8.13[c] | 44.50 ± 10.15[c] | 36.67 ± 15.91[c] | 45.17 ± 22.65[c] | 28.33 ± 8.89[c] | 26.17 ± 10.52[c] | 11.00 ± 4.00[c] |

Values presented are Mean ± SEM. [a]: Denotes statistically significant difference between A-B; [b]: Denotes statistically significant difference between B-C; [c]: Denotes statistically significant difference between B-D.

### Table S4. Protective effect of *Withania somnifera* against levamisole [Raw data pertaining to Figure 3(B)]

| | | 0 h | Day 1 | Day 2 | Day 3 | Day 4 | Day 5 |
|---|---|---|---|---|---|---|---|
| A | Worms incubated in M9 buffer | 111.11 ± 39.51 | 110.00 ± 14.10 | 85.00 ± 8.14 | 76.22 ± 17.51 | 67.33 ± 13.09 | 62.78 ± 15.54 |
| B | Levamisole pre-treated worms incubated in M9 buffer | 33.11 ± 28.05[a] | 66.00 ± 26.02[a] | 57.33 ± 26.49[a] | 42.11 ± 16.01[a] | 17.89 ± 6.83[a] | 18.78 ± 11.95[a] |
| C | Levamisole pre-treated worms incubated with *W. somnifera* | 33.44 ± 16.37 | 122.78 ± 38.41[b] | 120.00 ± 31.50[b] | 116.89 ± 33.52[b] | 99.56 ± 24.69[b] | 95.89 ± 15.43[b] |
| D | Levamisole pre-treated worms incubated with dopamine | 38.44 ± 13.35 | 100.50 ± 40.50 | 69.00 ± 31.21 | 123.89 ± 30.55[c] | 178.44 ± 40.73[c] | 370.78 ± 29.56[c] |

Values presented are Mean ± SEM. [a]: Denotes statistically significant difference between A-B; [b]: Denotes statistically significant difference between B-C; [c]: Denotes statistically significant difference between B-D.

### Table S5. Protective effect of *Withania somnifera* against $MnCl_2$ [Raw data pertaining to Figure 4(A)]

| | | 0 h | 15 min | 1 h | 3 h | 5 h | Day 1 | Day 2 | Day 3 | Day 4 | Day 5 |
|---|---|---|---|---|---|---|---|---|---|---|---|
| A | Worms incubated in M9 buffer | 79.00 ± 29.22 | 114.33 ± 24.11 | 104.00 ± 18.73 | 103.67 ± 13.26 | 121.33 ± 12.85 | 52.50 ± 18.92 | 46.33 ± 10.73 | 29.50 ± 14.29 | 23.67 ± 12.24 | 13.83 ± 7.99 |
| B | Worms challenged with $MnCl_2$ | 91.83 ± 13.57 | 39.67 ± 20.48[a] | 59.17 ± 6.68[a] | 48.00 ± 14.79[a] | 51.00 ± 6.57[a] | 1.83 ± 1.83[a] | 2.67 ± 3.61[a] | 4.33 ± 4.46[a] | 2.17 ± 3.25[a] | 0.17 ± 0.41[a] |
| C | *W. somnifera* pre-treated worms challenged with $MnCl_2$ | 95.33 ± 18.57 | 122.00 ± 13.33[b] | 133.17 ± 18.28[b] | 133.50 ± 17.85[b] | 129.50 ± 28.91[b] | 24.83 ± 11.53[b] | 22.33 ± 9.44[b] | 13.67 ± 3.14[b] | 8.50 ± 3.99[b] | 15.17 ± 5.00[b] |
| D | Dopamine pre-treated worms challenged with $MnCl_2$ | 82.67 ± 23.81 | 68.00 ± 15.11[c] | 107.17 ± 8.35[c] | 97.17 ± 23.15[c] | 108.17 ± 33.28[c] | 58.83 ± 18.39[c] | 36.83 ± 10.76[c] | 25.67 ± 9.37[c] | 29.50 ± 13.63[c] | 12.17 ± 6.74[c] |

Values presented are Mean ± SEM. [a]: Denotes statistically significant difference between A-B; [b]: Denotes statistically significant difference between B-C; [c]: Denotes statistically significant difference between B-D.

### Table S6. Protective effect of *Withania somnifera* against $MnCl_2$ [Raw data pertaining to Figure 4(B)]

| | | 0 h | 1 h | Day 1 | Day 2 | Day 3 | Day 4 | Day 5 |
|---|---|---|---|---|---|---|---|---|
| A | Worms incubated in M9 buffer | 111.11 ± 39.51 | 107.44 ± 38.50 | 110.00 ± 14.10 | 85.00 ± 8.14 | 76.22 ± 17.51 | 67.33 ± 13.09 | 62.78 ± 15.54 |
| B | $MnCl_2$ pre-treated worms incubated in M9 buffer | 26.22 ± 21.42[a] | 28.44 ± 20.39[a] | 33.78 ± 21.49[a] | 26.67 ± 18.92[a] | 21.00 ± 12.04[a] | 12.56 ± 8.73[a] | 12.00 ± 5.77[a] |
| C | $MnCl_2$ pre-treated worms incubated with *W. somnifera* | 31.22 ± 20.29 | 50.44 ± 11.36[b] | 80.11 ± 24.12[b] | 89.56 ± 23.72[b] | 98.11 ± 34.80[b] | 104.89 ± 33.33[b] | 98.89 ± 47.39[b] |
| D | $MnCl_2$ pre-treated worms incubated with cortisol | 38.56 ± 15.31 | 45.22 ± 18.57 | 66.00 ± 12.28[c] | 49.44 ± 14.27[c] | 39.44 ± 13.51[c] | 44.00 ± 16.85[c] | 26.89 ± 7.27[c] |
| E | MnCl2 pre-treated worms incubated with dopamine | 28.67 ± 10.25 | 43.00 ± 14.48 | 73.89 ± 11.25[d] | 47.56 ± 8.38[d] | 36.78 ± 6.02[d] | 79.11 ± 28.54[d] | 168.22 ± 78.73[d] |

Values presented are Mean ± SEM. [a]: Denotes statistically significant difference between A-B; [b]: Denotes statistically significant difference between B-C; [c]: Denotes statistically significant difference between B-D; [d]: Denotes statistically significant difference between B-E.
